# The clinical drug candidate anle138b binds predominantly to the central cavity in lipidic Aβ₄₀ fibrils and modulates fibril formation

**DOI:** 10.1101/2025.05.30.655790

**Authors:** Mookyoung Han, Benedikt Frieg, Dirk Matthes, Andrei Leonov, Sergey Ryazanov, Karin Giller, Evgeny Nimerovsky, Marianna Stampolaki, Kai Xue, Kerstin Overkamp, Christian Dienemann, Dietmar Riedel, Armin Giese, Stefan Becker, Bert L. de Groot, Gunnar F. Schröder, Loren B. Andreas, Christian Griesinger

## Abstract

Alzheimer’s disease is a specific neurodegenerative disorder, distinct from normal aging, with a growing unmet medical need. It is characterized by the accumulation of amyloid plaques in the brain, primarily consisting of amyloid beta (Aβ) fibrils. Therapeutic antibodies can slow down the disease, but are associated with potential severe side effects, motivating the development of small molecules to halt disease progression. This study investigates the interaction between the clinical drug candidate small molecule anle138b and lipidic Aβ_40_ fibrils of type 1 (L1). L1 fibrils were previously shown to closely resemble fibrils from Alzheimer’s patients^18^. Using high-resolution structural biology techniques, including cryo-electron microscopy (cryo-EM), nuclear magnetic resonance (NMR) spectroscopy enhanced by dynamic nuclear polarization (DNP), and molecular dynamics (MD) simulations, we find that anle138b selectively binds to a cavity within the fibril. This structural insight provides a deeper understanding of a potential drug-binding mechanism at the atomic level and may inform the development of novel therapies and diagnostic approaches. In addition, anle138b reduces fibril formation in the presence of lipids by approximately 75%. This may suggest a mechanistic connection to its previously reported activity in animal models of Alzheimer’s disease^15^.

## Introduction

Alzheimer’s disease (AD) is a chronic neurodegenerative disorder that causes brain damage and cognitive decline and remains a significant challenge with no currently approved cure. The accumulation of amyloid beta (Aβ) is considered a primary pathological hallmark of the disease^1–3^, and as the condition progresses, tau protein aggregation^4,5^ and hyperphosphorylation^6–8^ are observed. Aβ_40_ and Aβ_42_ are the two main Aβ isoforms, with the 2-residue longer peptide, Aβ_42_, being more prone to aggregation. Aggregates of both Aβ types and tau contribute to the formation of amyloid plaques and neurofibrillary tangles (NFTs) in the central nervous system. Thus, understanding the structural characteristics of Aβ aggregates is essential for developing effective therapeutic strategies against this debilitating disease.

Currently, antibody therapies targeting Aβ aggregates have shown promise in slowing cognitive decline associated with disease progression. However, these antibodies have also been reported to cause adverse effects when administered at high doses^4^. Therefore, recent research has increasingly focused on the discovery of small molecules that can inhibit amyloid-beta aggregation while effectively crossing the blood-brain barrier.

Building on previous research, multiple studies have used *in vitro* systems and patient-derived samples to investigate different Aβ fibril polymorphic structures using techniques such as cryo-electron microscopy (cryo-EM) and solid-state nuclear magnetic resonance (ssNMR). Among these polymorphs, the lipidic Aβ_40_ fibril^18–20^ of type 1, henceforth called L1 Aβ_40_ fibril, which is predominantly observed in the *in vitro* system developed in prior work, has been shown to resemble the pathological structures observed in the brains of patients.

The L1 Aβ_40_ fibrils consist of two protofilaments that share an identical fold and are symmetrically twisted together, forming a double-protofilament helical structure. These fibrils also maintain an approximate pseudo-2_1_ screw symmetry, which contributes to their structural stability. Each monomer is composed of three β-strands, with a long transition region (Ala21-Gly33, hereafter referred to as the loop region) located between the 2^nd^ β-strand (β2: Lys16-Phe20) and the 3^rd^ β-strand (β3: Leu34-Val36). A cavity is formed at the interface between the protofilaments arranged with 2 fold symmetry, with the loop regions from each monomer contributing to cavity formation. This cavity is centrally located along the fibril axis and is hereafter referred to as the central cavity (Supplementary Fig. 2b, c).

In particular, the β-sheet architecture and protofilament interactions of L1 Aβ_40_ fibrils are similar to those in patient-derived samples. These structural parallels indicate that L1 Aβ_40_ fibrils represent key features of AD pathology and thus serve as a valuable model for developing therapeutic strategies targeting Aβ aggregation. This study aims to use an *in vitro* system to simulate disease-relevant conditions and assess the effects of a clinically relevant small molecule candidate on Aβ_40_ fibrils, offering insights into disease-modifying mechanisms.

Recent advances in small molecule therapeutics have highlighted anle138b [3-(1,3-benzodioxol-5-yl)-5-(3-bromophenyl)-1*H*-pyrazole]^10^ as a promising drug candidate capable of modulating amyloid aggregation. Anle138b is a hydrophobic small molecule that can cross the blood-brain barrier, showing potential as a therapeutic agent for neurodegenerative diseases. It has demonstrated disease-modifying effects in various neurodegenerative disease models^10–17^, including an AD mouse model (APP/PS1), by reducing Aβ_42_ plaque formation and protecting mitochondrial integrity^15^. In addition, it targets multiple proteins implicated in neurodegenerative diseases, including tau (AD), α-synuclein (Parkinson’s, MSA), and prion protein (CJD)^10–17^. This broad spectrum of activity suggests that the mechanism by which anle138b inhibits Aβ_42_ accumulation may extend to Aβ_40_, making its effects on Aβ_40_ fibrils a significant research focus. Anle138b has completed Phase I clinical trials in healthy volunteers^16^ and Parkinson’s disease patients (NCT04685265) and is currently being evaluated in Phase II clinical trials for multiple system atrophy (MSA) (NCT06568237).

The present study investigates how anle138b binds to and affects the L1 Aβ_40_ fibrils. The structural and functional effects of anle138b treatment are analyzed under two conditions: pre-aggregation (pre-treatment) and post-fibril formation (post-treatment). By utilizing solid-state nuclear magnetic resonance (ssNMR), cryo-EM, and molecular dynamics (MD) simulations, we aim to gain a detailed understanding of the molecular interactions between anle138b and L1 Aβ_40_ fibrils.

## Results

### Anle138b inhibits L1 Aβ_40_ fibril formation

To assess the effect of anle138b on L1 Aβ_40_ fibril growth, Aβ_40_ monomers were aggregated in the presence of liposomes containing anle138b at three different **S**mall **M**olecule-to-**P**rotein molar **R**atios (SMPRs) of 0, 0.6, or 1.2. Thioflavin T (ThT) fluorescence and circular dichroism (CD) analyses revealed a concentration-dependent inhibition of fibril formation by anle138b (Fig. 1a, Supplementary Fig. 1). CD spectra showed β-sheet signatures under all conditions at the beginning of incubation after mixing Aβ_40_ with liposomes (Supplementary Fig. 1b), indicating similar initial secondary structures regardless of the presence of anle138b. However, after 48 hours of incubation, a strong β-sheet signal was retained in the control sample (SMPR 0), whereas β-sheet formation was significantly reduced in samples treated with anle138b (SMPRs 0.6 and 1.2). Notably, fibril formation was reduced most significantly at SMPR 1.2, indicating a strong dose dependence (Supplementary Fig. 1a, c). These results were further supported by two complementary approaches: measurement of 1D (^1^H)^15^N CP spectrum signal intensity and quantification of fibrils by negative stain EM, both of which indicated a dose-dependent inhibition of fibril formation by anle138b (Fig. 1b, c).

**Fig. 1:**
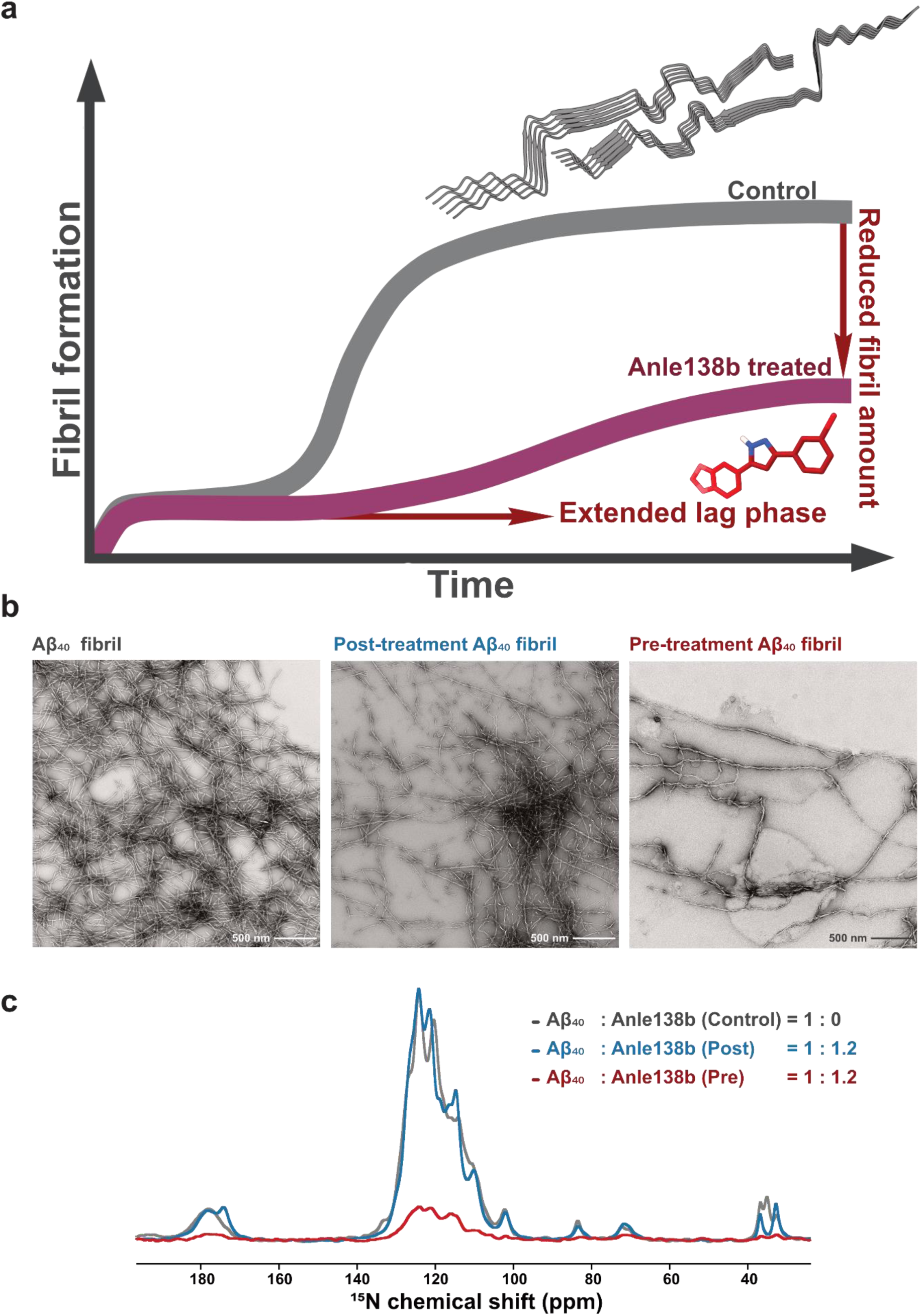
Anle138b modulates amyloid fibril formation: Comparison between pre-treatment and control conditions. **a.** Schematic representation of ThT fluorescence data showing fibril formation in the presence (red, pre-treatment condition) or absence (gray) of anle138b. The presence of anle138b extends the lag phase and reduces the final amount of fibrils. Full experimental data are provided in Supplementary Fig. 1. **b.** Negative stain electron microscopy (EM) images of Aβ_40_ fibrils. Left: Control fibrils formed without anle138b. Middle: Fibrils that were post-treated with anle138b (Aβ_40_: anle138b [SMPR] = 1:1.2). Right: Fibrils formed in the presence of anle138b (pre-treatment condition, [SMPR] = 1:1.2). **c.** 1D (^1^H)^15^N CP solid-state NMR spectra of Aβ_40_ fibrils. Gray, blue, and red spectra correspond to control, post-treatment, and pre-treatment Aβ_40_ fibrils. Spectral intensity in the pre-treatment sample is ∼25% of the control, consistent with ThT assay results and indicating reduced fibril formation in the presence of anle138b.

To evaluate the effect of anle138b at different stages of fibril formation, we compared three experimental conditions: 1) pre-treatment (SMPR 1.2, with anle138b added before initiating fibril formation), 2) post-treatment (SMPR 1.2, with anle138b applied after fibril formation was completed), and 3) control (fibril without anle138b) (Supplementary Fig. 2a). Fibril formation under the pre-treatment condition was reduced by approximately 75% compared to the control (Fig. 1c). In contrast, post-treatment resulted in no significant change in fibril quantity (Fig. 1c, Supplementary Fig. 3b). The observed reduction in ThT fluorescence intensity in the post-treatment sample (the fibril treated with anle138b after their formation) was therefore attributed to competitive binding between anle138b and ThT for fibril binding sites, rather than to an actual decrease in fibril quantity (Fig. 1b, c, Supplementary Fig. 3b). This explanation is consistent with previously reported interactions between small molecules and amyloid fibrils^21,22^.

In addition, CD spectroscopy and negative stain EM imaging confirmed that fibrils remained structurally unchanged after post-treatment with anle138b, even after 96 hours of incubation at 37°C (Supplementary Fig. 4). These findings indicate that mature fibrils retain their structural integrity in the presence of anle138b.

### Structural preservation and local effects of anle138b on L1 Aβ_40_ Fibrils

To determine whether the inhibition of fibril formation by anle138b is accompanied by structural changes, we analyzed the fibrils using cryo-EM and ssNMR. Consistent with previous findings, the ssNMR data^18^ indicate that L1 Aβ_40_ fibrils represent the predominant species of the control-fibril sample, and the L1 structure was reconstructed from the cryo-EM data for each preparation (control-fibril, post-treatment fibril, and pre-treatment fibril conditions). The cryo-EM analysis revealed no discernible structural differences between L1 Aβ_40_ fibrils in the presence or absence of anle138b, suggesting that the compound does not significantly alter the L1 Aβ_40_ fibril architecture (Fig. 2).

**Fig. 2:**
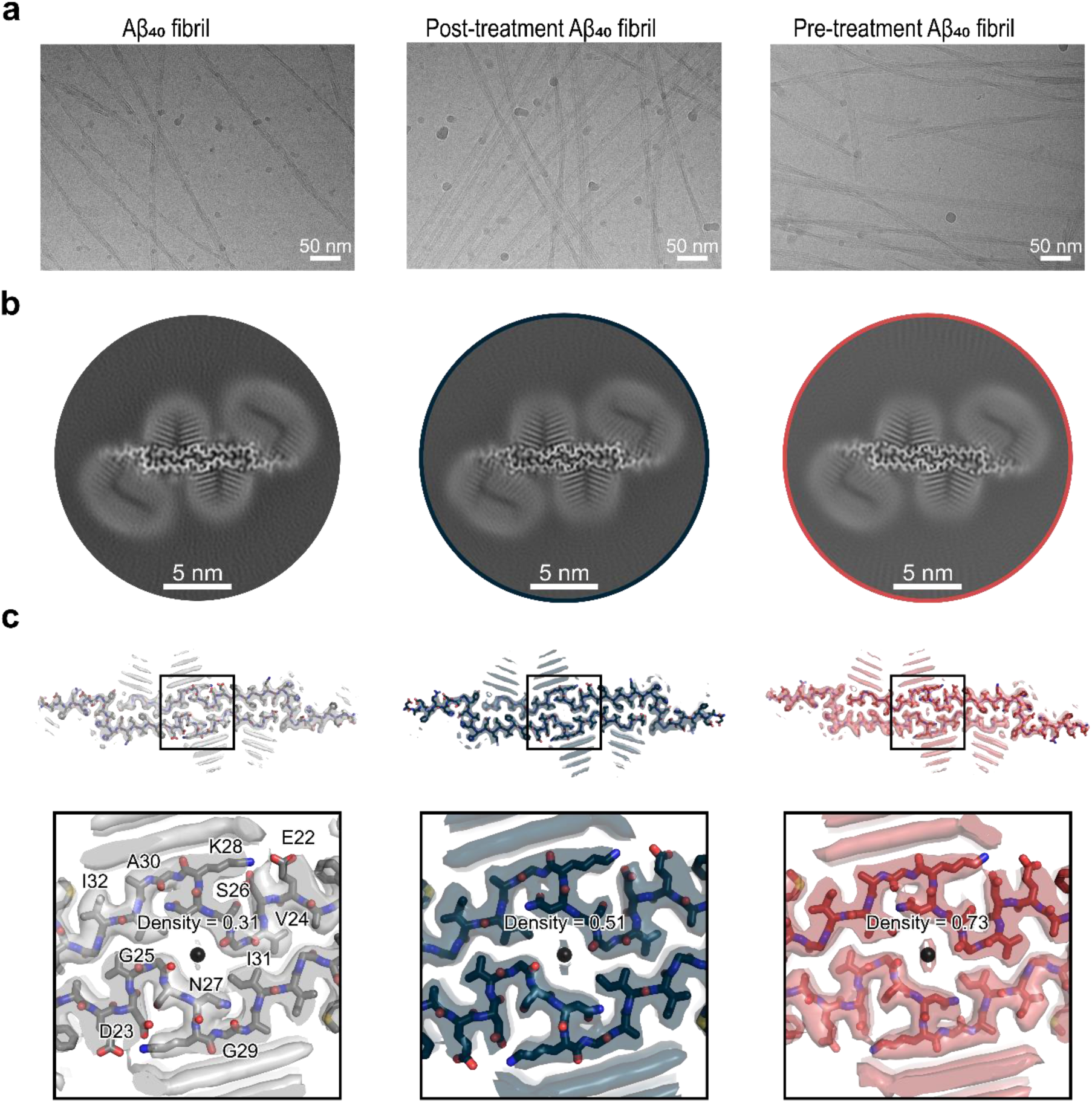
Cryo-EM maps reveal additional non-proteinaceous densities in L1 Aβ_40_ fibrils with anle138b. Lipidic Aβ_40_ fibril data in the absence of anle138b was previously published and is included here for comparison. **a**. Exemplary 20 Å low-pass filtered cryo-EM micrographs and 2D class averages for different fibril preparations: control condition (without anle138b), post-treatment condition, and pre-treatment condition. **b.** Cross-sections of the cryo-EM maps showing L1 Aβ_40_ fibrils under the three conditions. **c.** Overlay of a sharpened high-resolution cryo-EM map (transparent surface representation) and the atomic model (stick representation) of the fibril: control condition: gray (without anle138b), post-treatment condition: blue, and pre-treatment condition: red.

By contrast, ssNMR spectra revealed clear chemical shift perturbations (CSPs) in the presence of anle138b. In the 2D (H)NCA spectrum, substantial CSPs were detected for residues in the loop region between the second and third β-strand (Ala21–Gly33) (Supplementary Fig. 2b, c) of the L1 filament fold (Supplementary Fig. 5a). These spectral differences, relative to the untreated (control) fibrils, were observed following anle138b post-treatment of L1 Aβ_40_ fibrils (SMPR 1.2). Notably, the large CSP values of up to 10 ppm in ^15^N and ∼1.5 ppm in ^13^Cα suggest a direct interaction between anle138b and the fibril (Supplementary Fig. 5a).

Similar localized CSPs were observed under the pre-treatment condition (SMPR 1.2). This observation suggests that anle138b interacts with the loop region for both pre- and post-treatment conditions (Supplementary Fig. 5). Since these CSPs are observed across different parts of the L1 Aβ_40_ fibril structure, the data indicate that anle138b may bind to multiple regions of the fibril, including to the central cavity of the two protofilaments and also to exposed fibril surfaces. However, CSPs can also be the result of allosteric effects (*vide infra*).

In conclusion, the CSP results indicate that anle138b likely binds in the loop region (Ala21–Gly33), which includes the central cavity formed at the interface between the symmetry-related loop regions of the two protofilaments within the L1 Aβ_40_ fibril. Cryo-EM analysis indicates that the L1 Aβ_40_ fibril remains the predominant species in the presence of anle138b. The global fibril structure also remains unchanged.

### Anle138b binding in the central cavity of L1 Aβ_40_ fibrils

To directly assess binding locations, we employed amino acid-specific isotope labeling strategies and DNP-enhanced NMR. Chemical shift perturbations (CSPs) may be due to direct binding of anle138b to specific amino acids of L1 Aβ_40_ fibrils but could also be induced by allosteric structural changes. Therefore, DNP-enhanced NHHC experiments are required to localize the binding sites directly. For this purpose, the protein was labeled first uniformly with ^13^C and then with ^13^C, ^15^N-isoleucine or ^13^C, ^15^N-lysine, while anle138b was uniformly labeled with ^15^N at the central pyrazole group. The cross-peaks in the NHHC spectrum^64^ indicate close contact with the nuclei of the protein: the pyrazole ^15^N nuclei at 195 ppm are within approximately 5 Å of aliphatic ^13^C from the protein. These results provide direct evidence that anle138b binds to specific amino acids within the fibril (Fig. 3a, b), as detailed below.

**Fig. 3:**
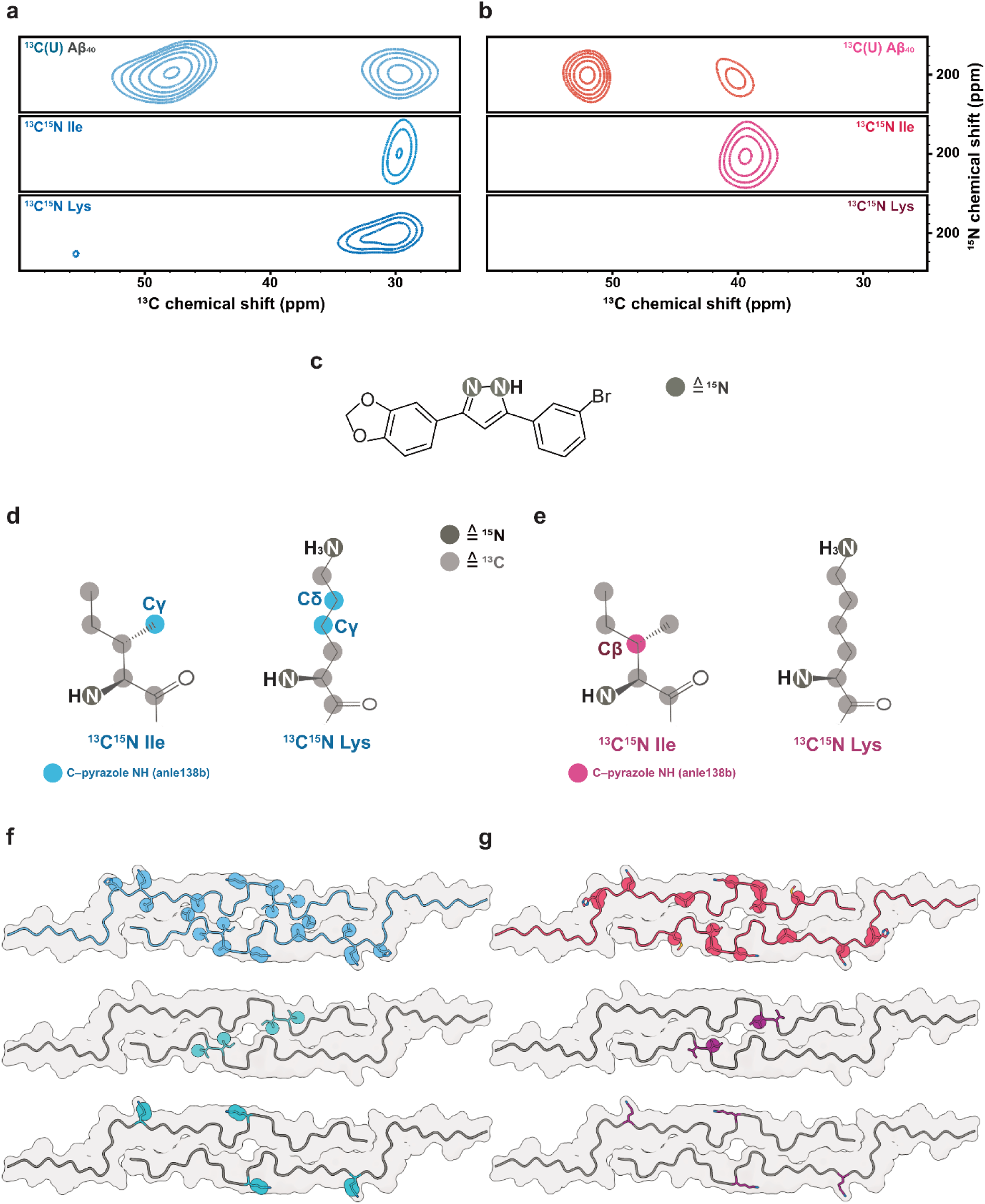
Interaction of anle138b with L1 Aβ_40_ fibrils: pre-treatment and post-treatment conditions. **a.** 2D NHHC spectra of L1 Aβ_40_ fibrils obtained after the addition of ^15^N-labeled anle138b under post-treatment conditions (Aβ_40_: anle138b [SMPR]=1:1.2, mixing time = 200 μs). The top panel shows uniformly ^13^C-labeled L1 Aβ_40_ fibrils, while the middle and bottom panels show spectra of fibrils selectively labeled with ^13^C, ^15^N-Ile31,32, and Lys16,28, respectively. Cross-peaks were observed between the ^15^N-labeled pyrazole group of anle138b and specific ^13^C nuclei within L1 Aβ_40_ fibrils. **b.** Same experimental setup as in **a**, except that anle138b was added before fibril formation (pre-treatment condition). **c.** Molecular structure of ^15^N-labeled anle138b, with labeling sites shown as dark gray circles. **d.** Chemical structures of labeled Ile and Lys residues under post-treatment conditions. ^13^C atoms identified in proximity to the pyrazole NH of anle138b are indicated in the blue circle. **e.** Same as in **d**, except for the pre-treatment condition. ^13^C atoms are marked in the red circle. **f.** Mapping of identified contact sites onto the L1 Aβ_40_ fibril structure under post-treatment conditions. Light blue spheres indicate ^13^C atoms from residues in the L1 Aβ_40_ fibril that are potential interaction sites for anle138b. Dark blue spheres in the middle and bottom models represent ^13^C atoms within Ile31,32 (middle) and Lys16,28 (bottom) that are in direct contact with anle138b. **g.** Same as in **f**, but for pre-treatment conditions. Red spheres indicate ^13^C atoms within contact residues.

Under the pre-treatment condition, a cross-peak appears between anle138b and the protein in the Cα region of the spectrum centered at 52 ppm, and a second peak is observed at about 40 ppm. The candidate residues that are consistent with these peaks include the Cα of His13, Ala21, and Ala30, and the sidechain resonances of Lys16, Lys28, Ile31, and Met35 (Fig. 3b, g). For a ^13^C, ^15^N-isoleucine–selectively labeled fibril, a cross-peak corresponding to Ile31 was identified as the interacting residue based on the Cβ chemical shift (Fig. 3b, g). In contrast, no cross-peaks were observed for a ^13^C, ^15^N-lysine–selectively labeled fibril, thereby excluding Lys16 and Lys28. Given that Ile31 points towards the central cavity and no contacts were detected with the lysine sidechains on the fibril surface, these data suggest that anle138b predominantly interacts with residues in the central cavity under the pre-treatment condition. The 52 ppm peak is consistent with contact with residue Ala30, which is next to Ile31.

In contrast, under the post-treatment condition, two prominent peaks corresponding to anle138b are observed at different positions, namely 49 ppm and 30 ppm (Fig. 3a, f) in the 2D NHHC spectrum of ^13^C-labeled protein. The Cα of residues such as Ala21, Ala30, Gly25, and Gly33 are consistent with the former peak, and the 30 ppm peak is consistent with the resonances of His14 (Cβ), Lys16, Lys28 (Cγ, Cδ), Ile31, Ile32 (Cγ1), Leu17, and Leu34 (Cδ). Leu17, Gly33, and Leu34 were excluded from consideration because of their buried location within the fibril. A DNP-enhanced 2D NHHC spectrum of isoleucine-labeled fibrils confirmed an interaction with Ile31 or Ile32. Lysine labeling also revealed a cross-peak at 30 ppm (Lys Cγ, Cδ), which, considering that these sidechains extend from the fibril, places anle138b on the fibril surface. These data confirm that anle138b interacts with surface-exposed residues such as Lys16, Lys28, and potentially Ile31 and/or Ile32. However, the small chemical shift difference (<1.35 ppm) between Ile31 Cγ1 and Ile32 Cγ1 resulted in peak overlap, preventing definitive assignment of the observed cross-peak to either residue. Therefore, binding to the central cavity cannot be definitively confirmed or excluded under the post-treatment condition.

Interestingly, the cryo-EM structures revealed an additional weak non-proteinaceous density within the fibril’s central cavity, going from 0.31 without anle138b to 0.51 in the post-treatment fibril condition to 0.73 in the pre-treatment fibril condition (Fig. 2c, d; Supplementary Fig. 6). Non-proteinaceous density (primarily lipids)^18^ is observed outside the fibrils, but there is no systematic density change depending on the presence of anle138b as for the central cavity.

### Post-treatment titration of the anle138b-L1 interaction

To assess the binding behavior under post-treatment fibril conditions, we characterized concentration-dependent interactions using ITC and NMR.

Isothermal titration calorimetry (ITC) revealed that anle138b binds to L1 Aβ_40_ fibrils with an exothermic and spontaneous profile (ΔH = –1.84 kcal/mol, ΔG = –8.45 kcal/mol). The dissociation constant (K_d_) was determined to be 0.64 μM, with a binding stoichiometry of ∼0.72 molecules per Aβ_40_ monomer (Fig. 4b, Supplementary Table 2). DLPG vesicles lacking anle138b produced no detectable heat change upon titration, indicating that the observed binding originated solely from interactions between anle138b and L1 Aβ_40_ fibrils (Supplementary Fig. 14b).

**Fig. 4:**
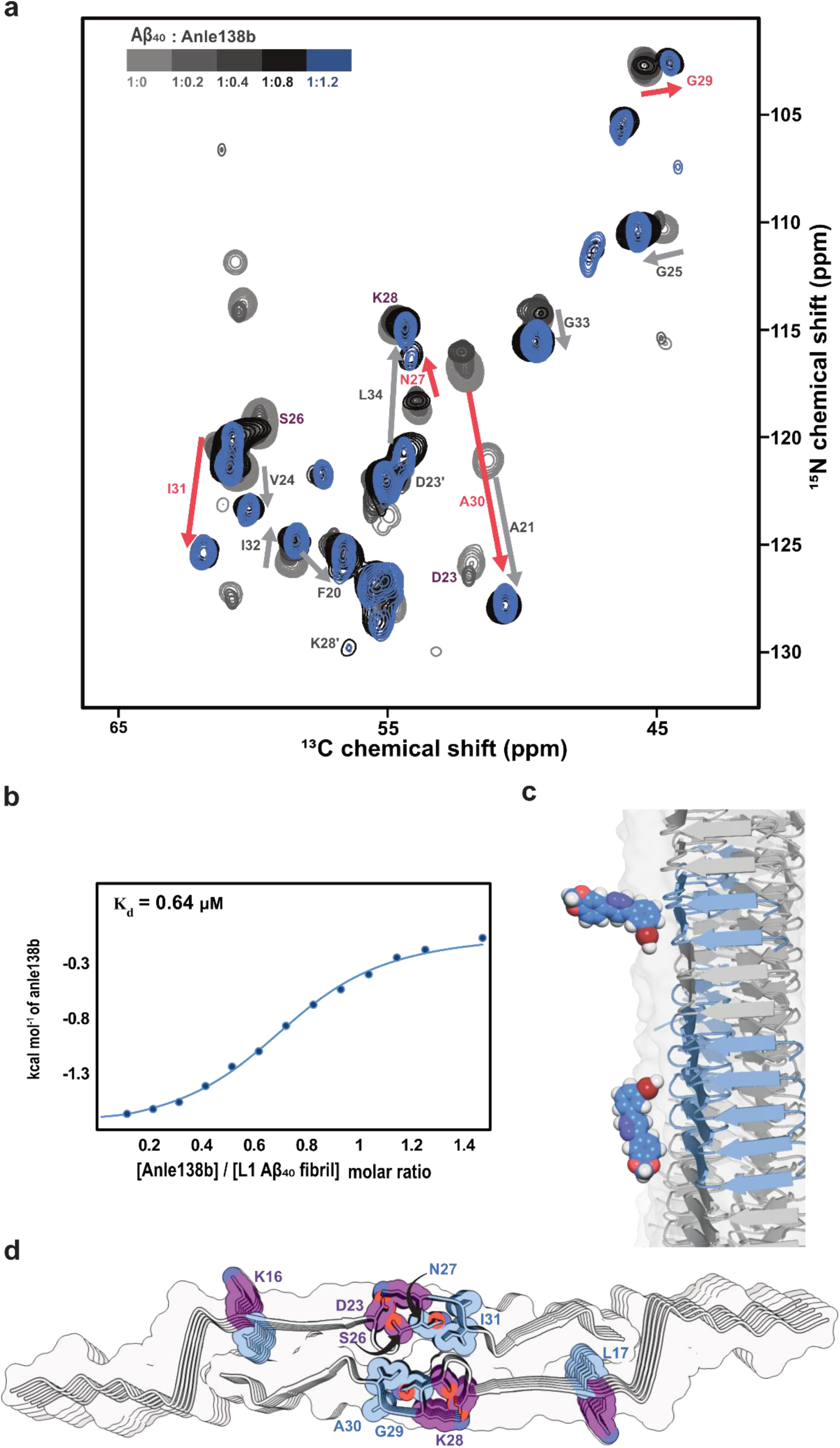
Titration of anle138b binding to L1 Aβ_40_ fibrils under post-treatment conditions. **a.** 3D (H)CANH solid-state NMR spectra of L1 Aβ_40_ fibrils titrated with increasing concentrations of anle138b (Aβ_40_: anle138b [SMPR] from 1:0 to 1.2). Chemical shift perturbations (CSPs) and signal intensity reductions are observed upon treatment with anle138b. Residues exhibiting both CSPs and signal attenuation are marked in pink arrows; those showing CSPs only are marked in gray arrows. Peaks for residues K16, D23, S26, and K28 (highlighted in purple) become undetectable at the highest anle138b concentration. **b.** ITC binding isotherm for anle138b binding to L1 Aβ_40_ fibrils, derived from integrated injection heats (solid blue circles) and fitted with a least-squares fit (blue line). The fitted binding isotherm yields a dissociation constant (K_d_) of 0.64 μM and a binding stoichiometry of ∼0.72 anle138b molecules per Aβ_40_ monomer. **c.** Structural modeling suggests two illustrative binding orientations of anle138b on the fibril surface: one approximately perpendicular (∼90°) and the other parallel (∼0°/180°) to the fibril axis, bridging up to 2 or 5 β-sheet layers, respectively. **d.** Perturbed residues are mapped onto the fibril structure. Blue indicates residues with both CSP ≥ 1.0 and intensity ratio (I_bound_/I_free_) ≤ 0.6. Purple indicates residues with undetectable signals at SMPR 1.2.

To resolve residue-specific changes and explore whether the binding stoichiometry observed in ITC correlates with the NMR spectra, we performed an NMR-based titration experiment using SMPRs of 0.2, 0.4, 0.8, and 1.2. The effects of binding were monitored via 3D (H)CANH spectra, analyzing CSPs and signal intensity changes (I_ratio). This revealed large chemical shift changes and slow exchange behavior in several residues (Fig. 4a).

Peaks associated with anle138b-bound fibrils emerged as early as SMPR 0.2 across the β2 strand, loop, and β3 regions, with residue- and region-specific differences in saturation behavior. Several residues exhibited both free and bound peaks between SMPR 0.2 and 0.8, with most sites saturating at 0.4 or 0.8. Notably, Gly29, Ala30, and Asn27 required SMPR 1.2 to reach saturation (Fig. 4a, Supplementary Fig. 11–13).

In more detail, in the **β2 region**, Leu17 showed early binding at SMPR 0.2 and saturated at 0.4, whereas Lys16 remained unaffected until signal loss at SMPR 1.2. In the **loop region**, multiple residues responded at low concentrations: Ala21, Gly25, and Gly33 reacted at 0.2, with Ala21 saturating at 0.4. Other loop residues (Gly29, Ala30, Ile31) responded at 0.4, while Asn27 showed delayed binding (slow exchange at 0.8; saturation at 1.2). Notably, Lys28 and Asp23 showed concentration-dependent signal disappearance at 0.4 and 0.8, respectively, and Ser26 was undetectable at 1.2. In the **β3 region**, Met35 responded early (0.2), and Leu34 entered slow exchange at 0.4 and saturated at 0.8. Together, these observations highlight region-specific differences in binding sensitivity, with the loop region displaying the broadest range of response behaviours.

Significant ^15^N chemical shift perturbations were observed for Ala30 (10.72 ppm), Ala21 (6.48 ppm), and Ile31 (4.65 ppm) in CSP-based titration experiments conducted under saturation conditions (SMPR 1.2) (Fig. 4a). These large shift changes in amide nitrogen signals can result from changes in the hydrogen bonding environment, and do not necessarily reflect a structural change in the protein ^67,72–75^. Indeed, upon anle138b binding, water-protein NOE contacts in the 3D H(H)NH NOE spectrum increased for Gly29, Ala30, Ile31, and Ile32, while decreasing for Gly25 and Leu34 (Supplementary Fig. 17).

The same spectrum also revealed that lipid interactions between β-strand residues (Leu17–Val18, Leu34–Val36) and DMPG acyl chains were disrupted, while new lipid contacts were formed with loop residues Ala21–Gly25 (Fig. 5). The disappearance of Asp23 and Lys28 signals indicates changes in the structure or dynamics of charged sidechains at the fibril surface near the loop region. Previous studies reported that Asp23 and Lys28 exist in two distinct populations (Asp23/Asp23′ and Lys28/Lys28′), of which only the Asp23–Lys28 pair forms a salt bridge^18^. In the 2D ^13^C^13^C-DARR spectra acquired under the SMPR 1.2 condition (mixing times = 20 and 200 ms), the peaks corresponding to the salt bridge conformation were absent, while only those corresponding to Asp23′ and Lys28′ were observed (Fig. 4a, Supplementary Fig. 8a, 15, 16). In addition to these residues, Ser26 also exhibited peak loss, and no cross-peaks were detected among Ser26(Cβ), Asp23(Cβ), Asn27(Cα/Cβ), and Lys28(Cε). Given that CP-based DARR spectra selectively detect rigid regions of the protein, these observations suggest enhanced flexibility or dynamic averaging in the loop region upon anle138b binding. At cryogenic temperature, differences in the chemical shifts of Lys28 Cγ and Cδ signals are evident: the room temperature assignments match pre-treatment fibrils, and these signals are absent in the post-treatment fibrils (Supplementary Fig. 10a). No major differences between pre- and post-treatment fibrils were observed for the isoleucine ^13^C resonances in the 2D ^13^C^13^C-DARR spectra acquired at cryogenic temperatures (Supplementary Fig. 10a). Note that the linewidths of ^13^C resonances at cryogenic temperature limit the detection of more subtle local differences. Taken together with cryo-EM analyses showing that all three fibril preparations— pre-, post-treatment, and control—share the same overall structure (Fig. 2), these findings suggest that the observed changes in the loop region primarily reflect enhanced local flexibility and water/lipid exposure rather than stable or static structural rearrangements.

**Fig. 5:**
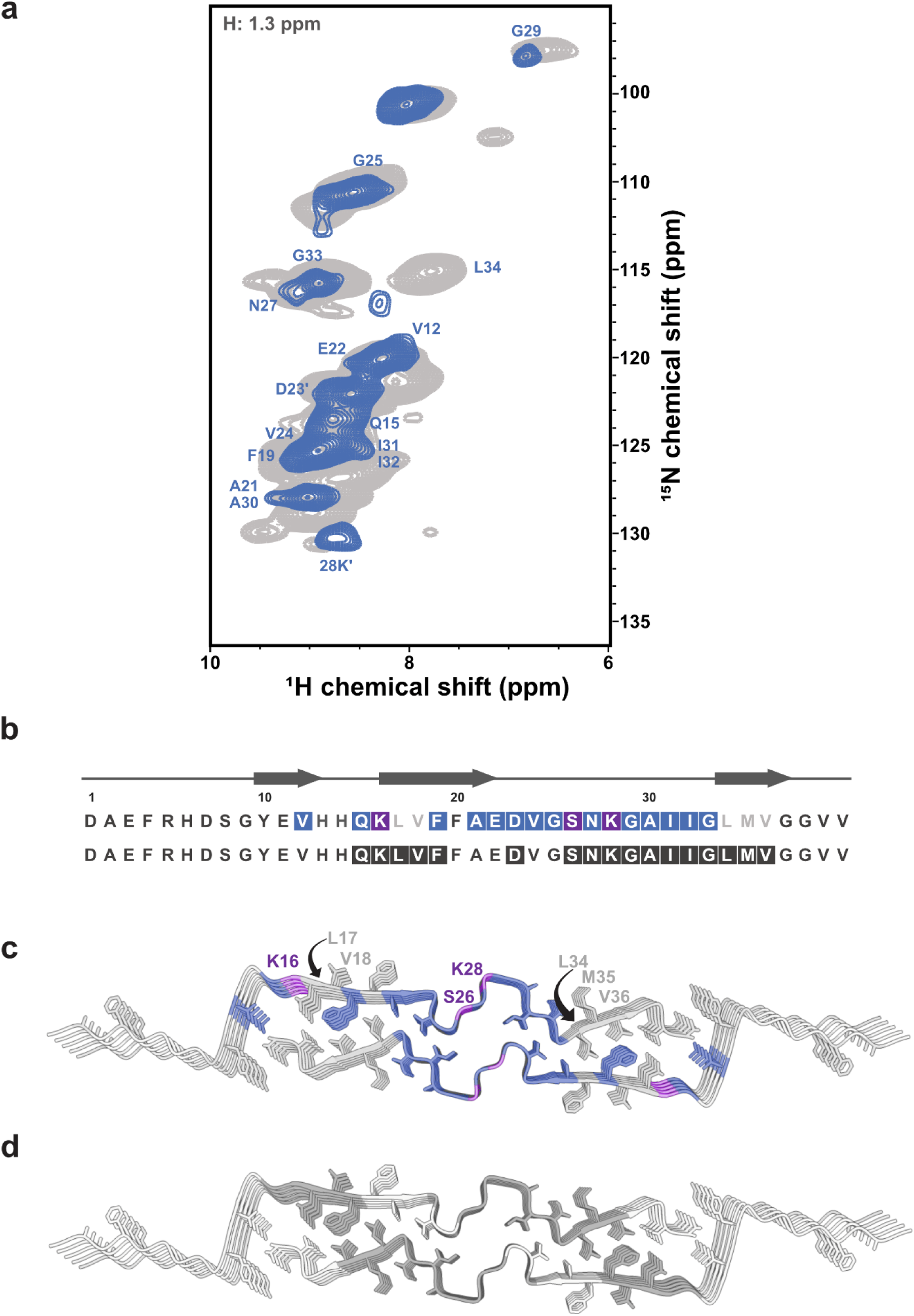
3D H(H)NH NOE analysis of lipid–fibril interactions in L1 Aβ_40_ fibrils after anle138b treatment (post-treatment condition) **a.** 2D (H)NH spectrum (gray) and 2D NH plane extracted at 1.3 ppm in the H dimension (lipid CH_2_ region) from a 3D H(H)NH NOE spectrum (blue, mixing time = 50ms), acquired on ^2^H, ^15^N, ^13^C-labeled L1 Aβ_40_ fibrils treated with anle138b (post-treatment; Aβ_40_: anle138b [SMPR] = 1:1.2). Blue cross-peaks mark backbone amide groups in proximity with lipid acyl chains. **b.** Amino acid sequence and secondary structure of L1 Aβ_40_ fibrils. Blue boxes (top) indicate residues interacting with lipid acyl chains (post-treatment condition, SMPR = 1:1.2). Light gray letters indicate loss of lipid contact post-treatment. Dark gray boxes (bottom) indicate lipid-interacting residues in the absence of anle138b. **c.** Structural mapping of lipid-contacting residues under post-treatment conditions. Blue color indicates interaction sites. Notably, hydrophobic residues (L17, V18, L34, M35, V36) are not involved in lipid interactions in the presence of anle138b. Purple labels mark residues that show signal loss after treatment. Hydrophobic side chains are shown as white sticks for reference (also in d). **d.** The same fibril model highlights lipid-interacting residues in the absence of anle138b (dark gray).

### Surface and central cavity binding modes explored via MD simulations

All-atom molecular dynamics (MD) simulations were employed to investigate the binding of anle138b to the L1 Aβ_40_ fibril structure observed by cryo-EM and NMR. We first considered the binding of anle138b, in the presence or absence of DMPG lipids, corresponding to an L1 Aβ_40_ fibril structure covered with and without surface-bound phospholipids (see Methods, Fig. 6). In both simulation sets, anle138b molecules established contacts with regions of the fibril surface, consistent with the NMR results of post-treatment fibrils (Fig. 6a-c). Notably, anle138b did not bind to the central cavity in either lipid-containing or lipid-free simulations. Lipid-free simulations show short distances (below 5 Å) between the pyrazole NH of anle138b and Cδ of residue Ile32, consistent with NMR data, while the lipid-containing simulations do not (Fig. 6b, c, Supplementary Fig.18a, b).

**Fig. 6:**
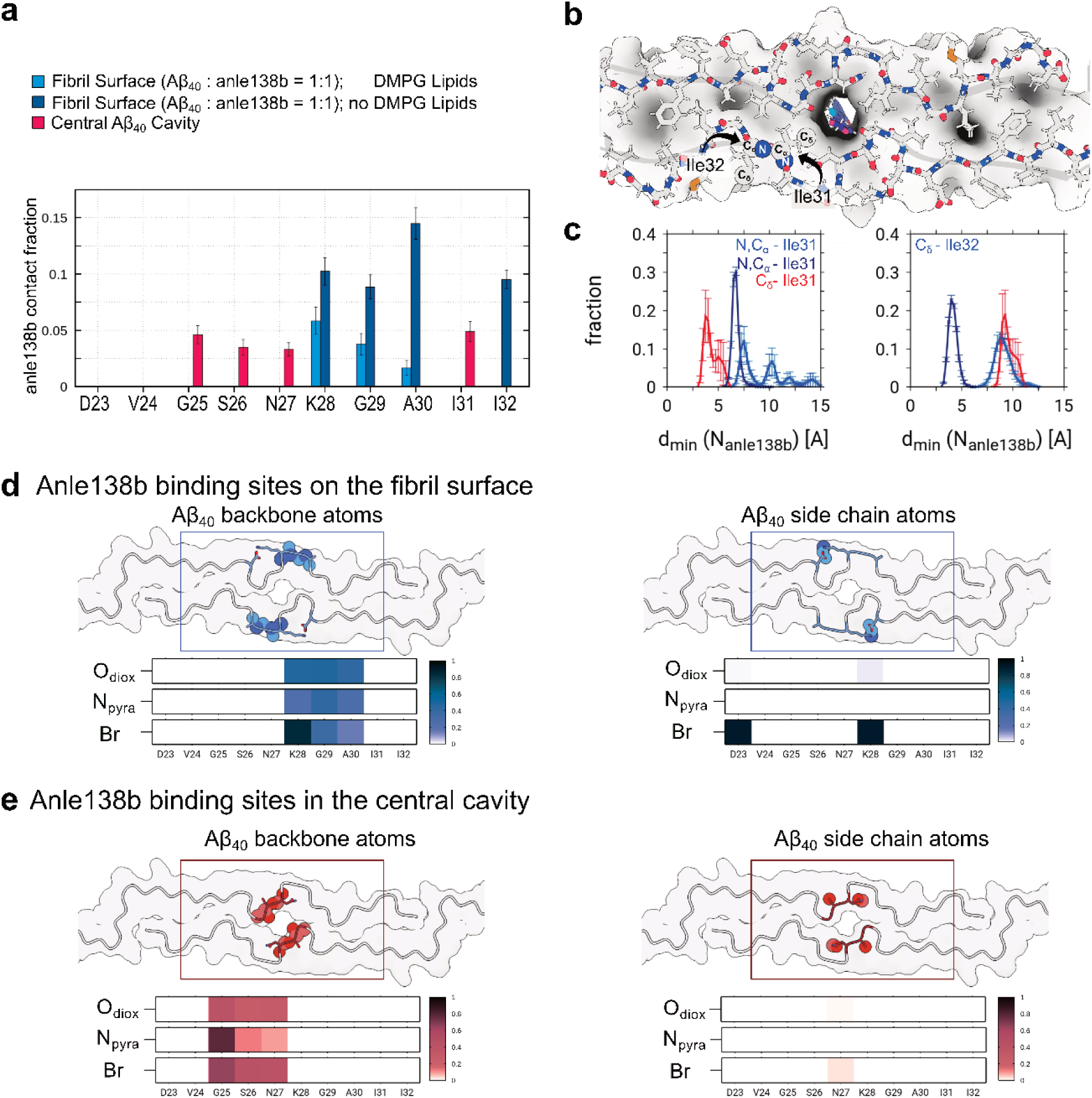
Structural determinants and contact sites of anle138b bound on the fibril surface and to the inner cavity of the loop region. **a.** Averaged anle138b contact fraction to the L1 Aβ_40_ fibril structure for MD simulations with anle138b molecules either bound to the surface (with DMPG lipids, blue and without DMPG lipids, dark blue) or internally bound to the loop region (red). Contacts are considered if protein N or C atoms to anle138b pyrazole nitrogen atoms are found within a cutoff distance of 0.5 nm. Data are presented as mean ± SEM (indicated by error bars). **b.** Representative MD simulation snapshot of L1 Aβ_40_ fibril loop region with internally bound anle138b viewed up close and down the long axis of the fibril. **c.** Distributions of distances between those N and/or C atoms of residues I31 and I32 that are minimal to anle138b pyrazole nitrogen atoms for the simulation sets representing either surface-bound (blue and dark blue) or internally bound (red) anle138b. Closest atomic contact sites of anle138b pyrazole nitrogen atoms are highlighted by spheres in **b** (annotated only for one of the protofilaments for simplicity). Renderings of L1 Aβ_40_ fibril loop region detailing contact sites accessible either to **d** Surface-bound or **e** Internally bound anle138b. Atoms from the backbone (left) and side chain (right) of the residues are highlighted separately. Renderings are accompanied by heat maps that report polar contacts between individual Aβ_40_ residues and polar moieties of anle138b, respectively. Scale bars indicate contact probabilities.

In a second set of simulations, we probed the ability of anle138b to spontaneously bind to the central cavity between the two protofilaments of the L1 Aβ_40_ fibril as observed under both pre-treatment fibril and post-treatment fibril conditions. To do so, the diffusion of anle138b in the simulation box was restricted to a cylindrical volume centered on the loop region by a flat-bottomed position restraint (see Methods, Supplementary Fig.19a, b). Indeed, anle138b can insert spontaneously in the central cavity of the loop region, showing contacts with Val24, Gly25, Ser26, Asn27, and Ile31 (Fig. 6a, b). During the microseconds-long MD trajectories, anle138b moved into the filament in eight out of ten simulation replicas, traversing up to 5 or 6 monomer layers into the central cavity. Once fully buried inside the fibril, the anle138b molecules are retained, as no unbinding events were observed (Supplementary Fig. 19a, b).

Interestingly, MD simulation models with deprotonated Lys28 and hence weakened side chain interactions between residues Asp23 and Lys28, allowed for a higher anle138b insertion rate (ten out of ten simulation replicas) and increased probability of deeper penetration into the central cavity of the fibril (Supplementary Fig. 19c). The disruption of the Asp23-Lys28 salt bridge led to an increase in RMSF values in the loop region, indicating enhanced structural flexibility (Supplementary Fig. 19d).

Anle138b molecules binding to the fibril surface and inside the central cavity exhibited contact patterns to residues Lys16, Lys28, Ile31, and Ile32, reproducing the two principal anle138b binding modes observed in the pre- and post-treatment fibril conditions by ssNMR experiments (Fig. 6a, c). Examination of the molecular determinants of anle138b binding revealed that the surface binding around the loop region occurred through contacts to the polar side chains of Asp23 and Lys28 (Fig. 6d, e). Specifically, the MD simulations revealed that the bromine (Br) atom of anle138b interacts with the ε-amino nitrogen (N) of Lys28 through a halogen bond-like contact^62^ (Fig. 6d, Supplementary Fig. 20a), whereas the central cavity binding was dominated by close polar interactions between anle138b and the exposed protein backbone atoms, particularly of Gly25 (Fig. 6e). The simultaneous interactions via the pyrazole and bromophenyl moieties allow for a favorable binding of anle138b to the fibril.

Compared to the distribution in the bulk lipid phase, the bromophenyl ring of surface-bound anle138b orients preferentially such that the bromine and pyrazole nitrogen atoms align on the same edge of the molecule (Supplementary Fig. 20b). Interestingly, the prominent binding to Asp23 and Lys28 with this ring orientation leads to a perpendicular alignment of anle138b with respect to the fibril axis and allows for close stacking of the ligands (Supplementary Fig. 20c). In contrast, anle138b binding to other regions of the L1 Aβ_40_ fibril surface, such as Lys16 or Gly37, as well as the central cavity, occurred with the molecules aligned parallel to the fibril axis.

In contrast to surface binding, the cavity binding mode showed no preferred bromophenyl ring orientation or anle138b alignment (bromophenyl moiety pointing down (0°) or up (180°) with respect to the fibril axis, Supplementary Fig. 18c, 20c, d). The observed anle138b binding mode characteristics were independent of the presence of DMPG lipids in the simulations, as well as the anle138b tautomer (Supplementary Fig. 20c, d).

## Discussion

### Disease relevance of the action of anle138b on L1 Aβ_40_ fibrils

This study builds on previous findings that L1 Aβ_40_ fibrils share structural similarity with Aβ_40_ fibrils amplified from amyloid seeds derived from the brain tissue of Alzheimer’s disease (AD) patients (PDB-ID: 6W0O)^19^. Similar fibril architectures have also been reported in Aβ_40_ fibrils amplified from vascular tissue from cerebral amyloid angiopathy (CAA) patients, including the familial Dutch variant (fCAA-Dutch, PDB-ID: 8FF3)^61^. These structural parallels and their pathological relevance support the use of L1 Aβ_40_ fibrils as a suitable model system for studying amyloid aggregation associated with AD and CAA.

Anle138b effectively inhibited Aβ_40_ fibril formation under pre-treatment conditions, consistent with previous findings of reduced Aβ_42_ deposition in the APP/PS1 mouse model^15^. Under these conditions, anle138b predominantly binds within a central cavity of the Aβ_40_ fibril, as revealed by ssNMR-detected contacts between anle138b and specific amino acids and corroborated by cryoEM density increase in the central cavity. This binding mode closely resembles that observed for α-synuclein^23,63^, where internal cavities arise from the repetitive fibril architecture and imperfect core packing. Similar cavities have been reported in some polymorphs of other amyloid fibrils^70^ and have been proposed to serve as favorable binding sites for small hydrophobic molecules^69,71^. In both L1 Aβ_40_^18^ and α-synuclein^24^ fibrils, such cavities are lined with glycine and isoleucine residues—glycine, due to its small size, may promote cavity formation, whereas isoleucine likely contributes to hydrophobic stabilization of the ligand.

In contrast, under post-treatment conditions, anle138b binds to the fibril surface without disrupting the overall fibril architecture, while still inducing localized structural perturbations. This binding mode resembles the interaction patterns of known amyloid-binding compounds such as Thioflavin-T, Congo Red^25,26^, and various PET tracers^27–30^, although high stoichiometric excess is used in most of these studies. NHHC spectra provided direct evidence of anle138b binding to surface-exposed side chains^30^, indicating preferential interaction at solvent-accessible regions of the fibril. 3D (H)CANH titration data showed region-specific binding dynamics at room temperature: in both β2 and β3 regions, some residues responded rapidly, while others exhibited slower exchange behavior. The loop region displayed a particularly distinct pattern—surface-facing sites saturated quickly, whereas inward-facing positions responded more gradually or only at higher ligand concentrations. Isothermal titration calorimetry (ITC) thermodynamically characterized the surface binding of anle138b to L1 Aβ_40_ fibrils, revealing exothermic binding with a dissociation constant (K_d_) of 0.64 μM and a stoichiometry of ∼0.72 molecules per monomer. This result was consistent with NMR titration data, which similarly indicated a stoichiometry of 0.7–0.8 molecules per monomer and revealed residue-specific differences in saturation behavior, suggesting that anle138b binds selectively to defined structural regions. The observed substoichiometric binding ratio can be explained with multiple binding sites for anle138b in surface-exposed regions of the fibril, rather than the central cavity. Structural modeling suggests that anle138b does not bind individually to each monomer but instead interacts with interfacial β-sheet surfaces that span multiple stacked monomer layers. When oriented parallel to the fibril axis, a single anle138b molecule may span up to five β-sheet layers or up to two layers when aligned perpendicular, providing a structural rationale for the ITC-derived binding stoichiometry.

The loop region of L1 Aβ_40_ fibrils (Ala21–Gly33) plays a critical role in forming the central cavity between the two protofilaments. DNP NHHC spectra reveal that, under post-treatment with anle138b, the compound binds to Cγ atoms of Ile and the Cγ and Cδ atoms of Lys28. These residues are located within the loop region and exposed on the fibril surface, indicating that anle138b interacts with the fibril at the surface under these conditions. Cryo-EM analysis confirms that the overall fibril architecture remains intact. However, the attenuation of loop residue signals in room-temperature CP spectra suggests that anle138b binding enhances the dynamic motion of the loop rather than inducing static structural changes. In particular, residues such as Ala21, Ala30, and Ile31 showed pronounced CSPs and concomitant changes in hydration, indicating that ligand binding perturbs their local physicochemical environment within the loop region. Furthermore, Gly25, located within the central cavity, shows a loss of hydration, consistent with displacement of water from the cavity upon anle138b binding. This interpretation is supported by the loss of the Asp23–Lys28 salt bridge on the loop surface, suggesting that anle138b binding correlates with increased loop flexibility. While solid-state NMR data indicate interactions at the fibril surface in the post-treatment fibrils, molecular dynamics simulations suggest an additional or complementary binding mode: upon deprotonation of Lys28, loop flexibility increases further, facilitating anle138b insertion into the central cavity.

Our results reveal a dual mode of action: inhibition of fibril formation during early aggregation and selective binding to mature fibrils without major structural disruption. From a thermodynamic perspective, it is intriguing that anle138b can stably associate with fibrils while simultaneously impeding their formation. Although the structural basis for the observed inhibition remains unresolved, one plausible explanation is that anle138b preferentially stabilizes early, non-fibrillar aggregates, thereby reducing the formation of mature fibrils. This mechanism is similar to that proposed for anle145c, a structurally related diphenyl-pyrazole (DPP) compound, which has been shown to inhibit hIAPP fibrillation and has been proposed to stabilize non-toxic oligomeric species through a thermodynamically driven process^67^. Moving forward, we aim to identify and structurally characterize these compound-stabilized early intermediates, which may hold the key to understanding the therapeutic mechanism of anle138b.

## Methods

### Protein expression and purification

Aβ_40_ fused to an N-terminal SUMO domain with a histidine tag was produced recombinantly, as previously described^18^. Briefly, uniformly ^13^C- and ^13^C,^15^N-labeled samples were expressed in a minimal medium supplemented with ^15^NH_4_Cl and ^13^C_6_-D-glucose. ^2^H,^13^C,^15^N-labeled Aβ_40_ was expressed in E. coli strain BL21(DE3) adapted to 100% ^2^H_2_O minimal medium supplemented with ^2^H_7_,^13^C_6_-β-D-glucose and ^15^N-ammonium chloride. For the production of amino acid-specific forward labeled protein (^13^C,^15^N-Lys, ^13^C,^15^N-Ile), the labeled amino acids were added to the minimal medium. The fusion protein was purified on nickel NTA resin (Macherey-Nagel), followed by digestion with tobacco etch virus (TEV) protease. Subsequently, the digestion mix was supplemented with 6 M guanidinium hydrochloride, and the SUMO domain and TEV protease were removed by another passage over the nickel resin. Final protein purification was performed by reversed phase HPLC on a C4 Vydac column. The purified peptide was finally lyophilized before use.

### Preparation of [1,2-^15^N_2_]-3-(1,3-benzodioxol-5-yl)-5-(3-bromophenyl)-1H-pyrazole

The synthesis and characterization of [1,2-^15^N_2_]-3-(1,3-benzodioxol-5-yl)-5-(3-bromophenyl)-1*H*-pyrazole (anle138b) have been previously reported in detail^23^, and the same procedure was followed in this study without any modification.

### L1 Aβ_40_ fibril preparation

The L1 Aβ_40_ fibril was prepared as described in a previous report¹⁸. Briefly, lipid films were generated from either 1,2-dimyristoyl-sn-glycero-3-phosphoglycerol (DMPG) or 1,2-dilauroyl-sn-glycero-3-phosphoglycerol (DLPG) by evaporating the solvent under a stream of N_2_, followed by lyophilization.

For fibril formation experiments, the DMPG film was hydrated and sonicated in 10 mM sodium phosphate buffer (pH 6.5) for 5 minutes to generate vesicles that promote Aβ_40_ aggregation. Under the pre-treatment fibril condition, anle138b was incorporated into DMPG during lipid film preparation, resulting in vesicles containing both lipid and anle138b. Control fibrils and pre-treatment fibrils were formed with DMPG liposome with or without anle138b.

The Aβ_40_ stock solution was diluted into buffer containing either DMPG liposomes or anle138b-loaded DMPG liposomes (small unilamellar vesicles, SUVs; diameter 30–100 nm) to yield final concentrations of 20 μM protein and 600 μM lipid. Samples were incubated at 37 °C for 1–2 days under quiescent conditions. Fibril formation was confirmed by circular dichroism (CD) spectroscopy and thioflavin-T (ThT) fluorescence.

For post-treatment fibril, DLPG lipid films containing anle138b were hydrated and sonicated for 15 minutes in 10 mM sodium phosphate buffer (pH 6.5), yielding lipid-based particles <10 nm in diameter to solubilize the hydrophobic compound.

Although the exact structural identity of DLPG particles remains unclear, with their small size (<10 nm) and physicochemical properties suggesting they are more likely micelles, non-bilayer structures, or ultrasmall unilamellar vesicles^82^, we refer to them as “vesicles” throughout this work for clarity. Control ITC experiments confirmed that DLPG vesicles alone exhibited no measurable binding enthalpy with L1 Aβ_40_ fibrils, supporting their role as a passive solubilizing vehicle for anle138b.

The anle138b-loaded DLPG vesicles were used to treat preformed L1 Aβ_40_ fibrils for subsequent ITC, DNP-NHHC, HCANH titration, and Cryo-EM experiments. With the exception of the ITC experiment, L1 Aβ_40_ fibrils were incubated at 37 °C for 1 hour to allow sufficient interaction prior to measurement.

### Aggregation assays

Aggregation assays were conducted under the same lipidic conditions used for fibril formation, either with DMPG liposomes (SUV) alone (control fibril condition) or DMPG liposomes (SUV) containing anle138b (SMPR 1:0.6, 1:1.2, pre-treatment fibril condition).

Aβ_40_ (20 µM) with liposome and thioflavin T (30 µM) were incubated in a non-binding black 96-well plate (Greiner Bio-One, item no. 655906) at 37 °C under quiescent conditions. Fluorescence signals were measured at 430 nm excitation (35 nm bandwidth) and 485 nm emission (20 nm bandwidth) using a Tecan Spark plate reader with double orbital shaking (1 min, 6 mm amplitude, 54 rpm, every 10 min). The manual gain was set to 40, with 30 flashes and an integration time of 40 µs. The z-position was calibrated using an empty well before each experiment. All measurements were recorded using Tecan Spark control software (v 2.2), and data were analyzed using Microsoft Excel.

### Solid-state NMR measurements

L1 Aβ_40_ fibrils were prepared using the same protocol with various isotopic labeling schemes, including uniformly ^2^H, ^13^C, ^15^N-labeled, ^13^C, ^15^N-labeled, ^13^C-labeled, and selectively ^13^C, ^15^N-labeled (at Ile or Lys residues). These fibrils were used for backbone assignments, lipid interaction studies, intermolecular contact analysis, and investigations involving anle138b.

The spectral similarity among the various L1 Aβ_40_ fibril preparations was confirmed using multiple techniques and isotopic labeling schemes, including 2D (H)NCA and 2D ^13^C– ^13^C DARR of ^13^C^15^N-labeled fibrils, 2D ^13^C–^13^C DARR of ^13^C-labeled fibrils, 2D ^13^C–^13^C DARR of selectively labelled fibrils, and 3D (H)CANH of ^2^H, ^13^C, ^15^N-labeled fibrils. For residue-specific backbone assignments, 2D (H)NH, 3D (H)CANH, and 3D (H)coCAcoNH experiments were performed. The 3D (H)CANH spectra were also acquired under different concentrations of anle138b for titration experiments.

All 2D ^13^C–^13^C DARR and 2D (H)NCA spectra were acquired on Bruker Avance III and NEO 850 MHz spectrometers equipped with 3.2 mm MAS HCN probes, operated at a magnetic field strength of 20.0 T, with MAS at 17 kHz and a sample temperature of 265 K. Titration experiments and 3D H(H)NH spectra (NOE mixing time = 50 ms) were recorded on ^2^H, ^13^C, ^15^N-labeled Aβ_40_ fibrils using a 1.3 mm MAS HCN probe spinning at 55 kHz at 235 K, on a Bruker Avance III HD 800 MHz spectrometer operating at a magnetic field of 18.8 T^31^. A 3D (H)CANH spectrum of post-treatment Aβ40 fibrils (SMPR 1:1.2) was acquired at 235 K on a Bruker Avance NEO 1200 MHz spectrometer (28.2 T) using a 1.3 mm MAS HCN probe at a MAS rate of 55 kHz. The hCANH spectrum acquired at 800 MHz validated the absence of spectral changes in the 1200 MHz spectrum.

Spectra were processed and analyzed using CcpNmr^32^, and peak intensities were determined by integration of cross peaks. Assignments to lipidic Aβ_40_ fibrils without anle138b have been reported previously^18^. Chemical shift perturbations were calculated as the average of the HN, NH, and Cα chemical shifts.

### DNP-enhanced ssNMR

TEMTriPol-1 was synthesized following established protocols^33^. The sample matrix, consisting of 45%^13^C-depleted d₈-glycerol and 4 mM TEMTriPol-1, was added to the fibril– lipid mixture and mixed to ensure homogeneous distribution.

All DNP experiments were performed on a Bruker Avance III HD 600 MHz spectrometer operating at 14.1 T, equipped with a 3.2 mm low-temperature (LT) MAS HCN probe. MAS frequencies used for each experiment are listed in Supplementary Table 1. Microwave irradiation at 395 GHz was generated using a gyrotron oscillator and transmitted via a corrugated waveguide. The sample temperature was maintained at ∼100 K using a Bruker second-generation liquid nitrogen cooling system.

2D NHHC experiments (mixing time = 200 µs) were conducted on ^13^C-labeled and selectively labeled lipidic Aβ_40_ fibrils in the presence of anle138b under both pre- and post-treatment fibril conditions. Additional 2D ^13^C–^13^C DARR and ^13^C–^13^C RFDR spectra were acquired on the same types of samples with anle138b applied in both conditions.

3D structural models were visualized using UCSF ChimeraX (v1.9)⁶⁶, and all graphs and plots were generated using Microsoft Excel.

### Affinity measurements

Isothermal titration calorimetry (ITC) was used to assess the binding affinity of anle138b to L1 Aβ_40_ fibrils in 10 mM sodium phosphate buffer (pH 6.5) at 25 °C, using a MicroCal PEAQ-ITC automated system (Malvern).

The experimental approach was based on established protocols for nanoparticle–protein interaction studies^79,80^ and adapted to systems involving drug-loaded vesicles^78,81^.

Anle138b (100 μM), prepared in DLPG vesicle (2 mM DLPG, particle diameter <10 nm), was titrated into the sample cell containing L1 Aβ_40_ fibrils (10 μM). The titration consisted of an initial 0.4 μL injection, followed by fourteen 2 μL injections, each lasting 4 seconds with 150-second intervals. As a control for dilution and background heat, DLPG vesicles without anle138b were titrated into the fibril solution using the same protocol. Control heat signals were subtracted from the raw thermogram to isolate the enthalpic contributions of anle138b binding. The ITC data were analysed using MicroCal PEAQ-ITC analysis software.

### Transmission electron microscopy

For negative staining, samples were applied to 400 mesh copper grids coated with glow-discharged carbon film. After staining with NanoVan (Nanoprobes Inc.), micrographs were taken at room temperature using a Talos L120C transmission electron microscope (Thermo Fisher Scientific).

### Cryo-EM sample preparation

When fibrils were formed with the same concentration of Aβ_40_ monomer for cryo-EM analysis, the post-treatment condition produced amounts of fibrils similar to the control condition, whereas the pre-treatment condition produced relatively fewer fibrils. To compensate for this difference, post-treatment fibrils were concentrated to 60 µM (consistent with the concentration used for the control condition in previous studies), while pre-treatment fibrils were concentrated to 720 µM, based on the initial Aβ_40_ monomer concentration. For this, 30 kDa Amicon concentrators (Ultra-500 μL) were used in a tabletop centrifuge at 12,000 g. After confirming similar fibril amounts based on CD signal intensity, cryo-EM grids were prepared using the concentrated pre- and post-treatment fibril samples.

### Cryo-EM grid preparation and imaging

The cryo-EM procedures, including grid preparation, data acquisition, and motion correction, were largely performed as described in our previous study^18^, with only the key parameters that differed in this study are detailed below.

Grids were flash-frozen in liquid ethane using a Mark IV Vitrobot (Thermo Fisher) operated at 95% relative humidity. Data acquisition was carried out in EFTEM mode using a Quantum LS (Gatan) energy filter with a slit width of 20 eV. A total of 7,311 and 21,576 movies were collected using SerialEM^76^ for pre- and post-treatment condition samples, respectively. Each movie was recorded over 40 frames with a total accumulated dose of ∼40.5 e⁻/Å^2^. The defocus range was set from −0.7 to −2.4 μm. Motion correction and dose-weighting were performed on-the-fly using Warp^77^.

### Helical reconstruction of Aβ_40_ fibrils

Aβ_40_ fibrils in complex with anle138b were reconstructed using RELION-3.1^34^, following the helical reconstruction scheme^35^. First, the estimation of contrast transfer function parameters for each motion-corrected micrograph was performed using CTFFIND4^36^. Next, filament picking was done using crYOLO^37^.

The reconstruction procedure was adapted from the original L1 Aβ_40_ fibril. Hence, for 2D classification, we extracted particle segments using a box size of 600 pix (1.05 Å/pix) downscaled to 200 pix (3.15 Å/pix) and an inter-box distance of 13 pix. For 3D classification, the classified segments post-2D classification were (re-)extracted using a box size of 250 pix (1.05 Å/pix) and without downscaling. Starting from a featureless cylinder filtered to 60 Å, several rounds of refinements were performed while progressively increasing the reference model’s resolution. The helical rise was initially set to 4.75 Å, and the twist was estimated from the micrographs. Once the β-strands were separated along the helical axis, we optimized the helical parameters (final parameters are reported in Tab. S4). We then performed a gold-standard 3D auto-refinement, followed by standard RELION post-processing with a soft-edged solvent mask that includes the central 10 % of the box height yielded post-processed maps (B-factors are reported in Tab. S3). The resolution was estimated from the value of the FSC curve for two independently refined half-maps at 0.143 (Fig. S7). The optimized helical geometry was then applied to the post-processed maps yielding the final maps used for model building.

### Atomic model building and refinement

For the Aβ_40_ fibril, one protein chain was extracted from PDB-ID 8ovk. Subsequent refinement in real space was conducted using PHENIX^3,38^ and Coot^39^ in an iterative manner. The resulting models were validated with MolProbity^40^, and details about the atomic models are described in Tab. S4.

### MD simulations

#### Simulation protocol

The GROMACS 2023 simulation software package^41,42^ was used to set up and carry out the MD simulations. Settings for production runs were chosen as follows: The long-range electrostatic interactions were treated using the Particle Mesh Ewald (PME) method^43,44^. Bonds in protein and lipid molecules were constrained using the P-LINCS^45^ algorithm. Water molecules were constrained using SETTLE^46^ algorithm. Neighbor lists were updated with the Verlet list scheme^42,47^. For production runs, the simulated systems were kept at a temperature of 300 K by applying the velocity-rescaling^48^ algorithm. Initial velocities for the production runs were taken according to the Maxwell-Boltzmann distribution at 300 K. The pressure was held constant by using the Parrinello-Rahman barostat^49^. The integration time step was set to 2 fs. The neighbor lists for non-bonded interactions were updated every 20 steps. Real-space electrostatic interactions were truncated at 1.2 nm. The van der Waals interactions were switched off between 1.0 to 1.2 nm and short-range electrostatic interactions were cut-off at 1.2 nm. All simulations were carried out using periodic boundary conditions for the simulation box and utilized the CHARMM36m^50,51^ protein force field together with the CHARMM-modified^52^ TIP3P water model and the CHARMM36 lipid parameters^53^. The openbabel web server^54^ was used to create the input files for topology and parameter preparation of anle138b molecules with the CHARMM General Force Field (CGenFF 4.6)^55–58^. The topologies were converted to GROMACS format using the cgenff charmm2gmx.py script (http://mackerell.umaryland.edu/charmm_ff.shtml#gromacs). All production simulations were preceded by a multi-step energy minimization and thermalization of the simulation systems. Atomic coordinates for analysis were stored every 100 ps.

### Setup of full-length L1 Aβ_40_ model

To investigate lipid and small molecule binding to different parts of the lipidic polymorph L1 Aβ_40_ fibrils located by ssNMR measurements, we performed MD simulations of an Aβ_40_ fibril model in the presence of DMPG lipids. The cryo-EM structure of L1 Aβ_40_ fibrils (PDB-ID: 8ovk) was used to build a fibril model with two protofilaments, each composed of 25 helically arranged peptide chains. If not stated otherwise, the titratable groups of the protein structure were protonated according to their standard protonation states at pH 7. The L1 Aβ_40_ fibril structure was placed in a rectangular solvent box with the distance between the solute and the box edge of at least 2.5 nm.

Based on this principal simulation system, two setups were built by adding the small molecule anle138b (representing both possible pyrazole nitrogen protonation states in a 1:1 ratio):

a. 50 anle138b molecules were placed randomly in the solvent around the L1 Aβ_40_ model structure. In the next step, 700 DMPG molecules were placed randomly in the solvent around the L1 model structure. The final Aβ_40_ monomer to DMPG lipid to anle138b ratios were 1:14:1. Sodium and chloride ions were added to yield an ionic strength of 150 mM and to neutralize the net system charge. The final setup of this L1 Aβ_40_ model contained 408,900 atoms, including 99,900 water molecules. Ten independent NPT production simulations, each 1.0 μs long, were carried out for this simulation system.
b. 50 anle138b molecules were placed randomly in the solvent around the L1 Aβ_40_ model structure. Sodium and chloride ions were added to yield an ionic strength of 150 mM and to neutralize the net system charge. The final setup of this L1 Aβ_40_ model contained 448,600 atoms, including 138,700 water molecules. Ten independent NPT production simulations, each 0.25 μs long, were carried out for this simulation system.

### Setup of truncated fibril model systems to study putative internal anle138b binding site

To investigate anle138b, the binding to the loop region of Aβ_40_ fibrils, and the dynamics of internally bound anle138b, we performed MD simulations on a shorter, restrained substructure. Instead of simulating the whole fibril model, only peptide chain parts near the suspected internal small molecule binding sites were included (PDB-ID: 8OVK, residues 15-39). The truncated simulation model of the L1 Aβ_40_ fibrils was composed of 30 helically arranged peptide chains per protofilament. The truncated simulation model of L1 Aβ_40_ fibrils were placed in a rectangular solvent box with a distance between the solute and the box edge of at least 1.5 nm. The titratable groups of the protein structure were protonated according to their standard protonation states at pH 7. Acetyl and N-methyl groups, respectively. Counterions (Na+, Cl−) were added to yield an ionic strength of 150 mM and to neutralize the net system charge.

For the initial configuration, one anle138b molecule was placed outside and above the central cavity of the loop region of L1 Aβ_40_ fibril, such that the long axis of the small molecule aligned with the fibril axis. Five independent NPT production simulations per possible pyrazole nitrogen protonation state of anle138b, each 5.0 μs long, were carried out for this simulation system.

An otherwise identical simulation with L1 Aβ_40_ fibrils featuring deprotonated lysine 28 side chains was performed for five independent NPT production simulations per possible pyrazole nitrogen protonation state of anle138b, each 5.0 μs long.

A flat-bottomed position restraint was used to keep the diffusion of the small molecule in the simulation box restricted to a cylindrical volume centered on the binding site of interest (radius 1 nm). This approach reduces computational cost significantly and increases the sampling efficiency of binding events to the cavity of the L1 Aβ_40_ fibril polymorph.

During all production MD simulations, the initial atomic coordinates of the Cα atoms of protein residues Asp1-Lys16 and Val36-Val40 were restrained using a harmonic potential with a force constant of 1000 kJ mol^−1^ nm^−2^ to ensure that the fibril models were stable, preserving the initial fold of the N- and C-terminal regions of the L1 Aβ_40_ fibril as resolved by cryo-EM.

### Reference simulations of anle138b in lipids

Anle138b conformations without *β*-sheet aggregates were set up and simulated as follows: A simulation box consisting of a solvated DMPG membrane patch composed of 76 lipid and 5266 water molecules and one anle138b ligand comprising 24,451 total atoms were simulated 6 x 1000 ns.

### Analysis and visualization

The MD simulation trajectories were preprocessed and analyzed by tools provided by the GROMACS 2023 simulation software package^41,42^, as well as custom-made bash, awk, and python scripts. Renderings of atomic coordinates were carried out with the PyMOL molecular visualization software^59^ [http://www.pymol.org/pymol], and the 3D structural model was visualized using UCSF ChimeraX version 1.9.^66^. Graphs and figures were created using Gnuplot 5.4 and Inkscape 1.3.

### Contact analysis

Aβ_40_ fibril residues are quantified to be in contact with anle138b using the g_contacts and gmx mindist programs^60^. In accordance with the type of NMR experiments and the isotope labeling scheme, the following contact criteria were chosen a) If the 1,2-N atoms of the anle138b molecule were found within a cut-off of 0.5 nm from any carbon or nitrogen atom of a protein residue, a contact was considered formed. b) In addition, a close interaction was, considered formed if all polar heavy atoms of anle138b (N, O, Br) molecules were found within a cut-off of 0.4 nm from all polar backbone or side chain heavy atoms of a protein residue. Contacts were averaged over time and over the individual trajectories of each simulation set. If not explicitly stated, only contacts to the core β-strand layers (neglecting the two β-strands on both protofilament edges) were considered for analysis.

### Data Availability

NMR: All thermodynamic data, titration curves, and NMR spectra have been deposited in the Biological Magnetic Resonance Data Bank under entry ID 53129. Cryo-EM: Cryo-EM density maps have been deposited in the Electron Microscopy Data Bank (EMDB) under accession numbers: EMD-53882 (pre-treatment fibril) and EMD-53880 (post-treatment fibril). The corresponding atomic models have been deposited in the Protein Data Bank (PDB) under accession codes 9RAX (pre-treatment) and 9RAW (post-treatment). MD simulation: Ligand topology and parameters, initial coordinates and simulation input files, and a coordinate file of the final output are provided through the Edmond data repository at [https://doi.org/10.17617/3.NRYUVQ].

## Acknowledgments

This work was supported by the Max Planck Society (to CG) and the German Research Foundation (DFG) in the framework of the German Excellence Strategy-EXC 2067/1-390729940 (to CG) and the Emmy Noether Program to LBA (project number: 397022504). BF and GFS are grateful for the computational support and infrastructure provided by the Zentrum für Information und Medientechnologie (ZIM) at the Heinrich-Heine-Universität Düsseldorf and for the computing time supplied by the Forschungszentrum Jülich on the supercomputer JURECA-DC at the Jülich Supercomputing Center (JSC). We would like to thank Prof. Markus Zweckstetter for providing access to the Tecan Spark plate reader and the MicroCal PEAQ-ITC automated system.

## Author contribution

GFS, LBA, B.L.dG., and CG designed the project. MH, BF, GFS, LBA, and CG managed the project. KG, KO, and SB performed the protein expression and purification. MH prepared the fibril samples. MH, EN, and MS performed the ssNMR experiments. CD prepared the cryo-EM grids and collected the cryo-EM images. BF processed the cryo-EM images, reconstructed the fibril structures, built the atomic models, and DM carried out MD simulations. MH, BF, and DM visualized the results. GFS, LBA, B.L.dG., and CG supervised the project. MH, BF, DM, B.L.dG., GFS, LBA, and CG wrote the original manuscript. All authors revised and edited the manuscript.

## Additional Information

Supplementary Information is available for this paper. Correspondence and requests for materials should be addressed.

